# Dynamic of Lotka-Volterra model for tumor-host systems under constant or periodic perturbation: implications for the therapy of cancer

**DOI:** 10.1101/2025.03.13.642966

**Authors:** Rolando Placeres Jiménez, Luis E. Bergues Cabrales, Juan I. Montijano

## Abstract

In this paper, the interaction tumor-host is described by a Lotka-Volterra model. The critical parameters that define the possible dynamical regimes are determined using linear stability analysis. The effects of constant and periodic perturbations are discussed, as well as their implications in clinics. The treatment dose required to lead the system to a desired state is obtained. It is also shown that aggressive tumors evolve to a limit cycle when the host is under the action of treatment applied periodically with a low frequency. A transition to a non-chaotic attractor occurs for higher frequencies. This transition tends to contract with the increment of the frequency of this external periodic perturbation. It is not detected chaotic behavior, even for higher values of both the strength and the frequency of the perturbation because the maximum Lyapunov exponent remains negative. These results may suggest that although aggressive tumors cannot be completely eliminated by conventional anticancer therapies, they might be controlled using external periodic therapies when only the host is perturbed.

## Introduction

The cancer is a leading cause of death worldwide despite the significant advances in the molecular and cellular biologies, medicine and biophysics. The effectiveness of many anticancer therapies are limited and a complete cure for cancer has not been achieved [1]. Anticancer therapies used in experiments have remained until present largely empirical, based on trial and error [2, 3]. The integration of mathematical modeling with experimental oncology is a necessary step for a better understanding of the genesis, growth, progression, invasion and metastasis of cancer, as well as individualized/personalized anticancer therapy targeting cancer hosts, whether patients (in clinical studies) or laboratory animals (in preclinical studies) [2, 3].

Mathematical models of population ecology provide a correct theoretical framework to describe the dynamics of interaction between malignant and normal cells [4–9]. Population models may be a useful theoretical tool to understand the tumor-host dynamic under the action of different sort of existing anticancer therapies, whether oncoespecific (chemotherapy and radiotherapy) or under study (biological (e.g., immunotherapy) and physical (e.g., electrochemical ablation, irreversible electroporation) therapies).

Population models may contribute to improve the effectiveness of anticancer therapy types. For this, the Lotka-Volterra (LV) model has been successfully used [4–9]. A distinctive feature of LV model is that population growth is limited and the complete remission may be induced in some tumor types using radiotherapy while accounting for information from both the tumor and host states [9]. Nevertheless, this anticancer therapy fails in advanced-stage tumors.

Many authors have studied the effects of periodic coefficients and different external perturbations in the LV model [10–12]. These studies are based on the fact that the original LV model without the logistic term has an oscillating behavior [13–15]. The competitive LV model is successfully applied to several problems in ecology [16–23]. Cushing [20, 21] and Gopalsamy [22] demonstrate that two competing species that are affected by an external periodic perturbation may coexist without each being eliminated by the other, avoiding extinction. A similar behavior has been observed in immune mathematical models [24–26] under the action of an external periodic perturbation. These aspects have allowed some researchers to suggest the use of external periodic perturbations as possible as therapeutic strategies to control cancer [25].

In the first section, the dynamic of the LV model is reviewed based on linear stability analysis from which we determine the set of parameters that define each dynamical regime, and the non existence of periodic orbits is proved, which means that the tumor-host tends always to one of the equilibrium point (destruction of the host, destruction of the tumor or coexistence of a fixed amount of tumor and host cells). In the second section, the effect of an external perturbation (cancer therapy) is studied and its implications in clinics are discussed Three different types of therapies are analyzed: Treatment applied continuosly, having an effect proportional to the cells population, treatment applied continuosly, having an effect independent of the cells population, and treatment applied periodically, having an effect independent of the cell populations. To help in this analysis, the LV equations are decoupled, which allows studying the dynamic of the system as the motion of a particle in a force field. Then, the characteristic frequency of oscillation around stable fixed points is defined. The dynamic of the perturbed system is studied using the phase space portraits and it is found the value of the dose of the applied treatment required to lead the tumor-host system to the different regimes. Moreover, the maximum Lyapunov exponent. is computed in the periodic ally applied treatment to show the non chaotic behaviour of the system. In the last section, we summarize the main results of this study and new insights derived of it that will be developed in a further study.

In experimental oncology, the action of internal (or endogenous) perturbations in the host and/or external (or exogenous) perturbations to the host may change tumor-host dynamics, tumor growth kinetics, and the biological characteristics and electrical properties of both tissues. Furthermore, it is common to include primarily tumor-directed anticancer therapies in studies of tumor-host dynamics. Nevertheless, it is no explored whether the host alone may govern this dynamics in the absence of external perturbations or when the host is only perturbed with them without the need to directly damage the tumor. That are why, both types of perturbations are addressed in separate sections of this study: Sections 2 and 3 (withouth external pertubation action), and Section 4 (with external pertubation action).

### The model and linear stability analysis

We consider two interacting species competing for space and resources, i.e., the cancer cells population *x* and the host cells population *y*. The evolution of this competence between these two population types may be modelled by the LV equations [4–9], given by

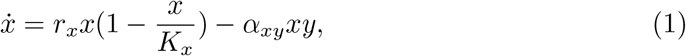

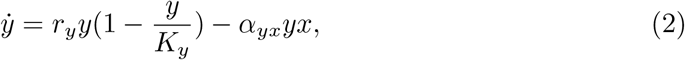

where the parameters *K*_*x*_ and *K*_*y*_ represent the carrying capacity of the cancer cells and host cells, respectively. The parameter *r*_*x*_ denotes the growth rate of cancer cells population, whereas *r*_*y*_ represents the growth rate of host cells population. The parameter *α*_*xy*_ is a measure of the interaction established by the host cells population in the presence of cancer cells population and *α*_*yx*_ is a measure of the interaction established by cancer cells population with the host cells population. All these parameters are assumed as non-negative constants and we restrict the analysis to the positive quadrant.

Using the following transformation of variables:

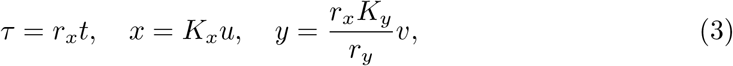

the equations (1) and (2) reduce to the form

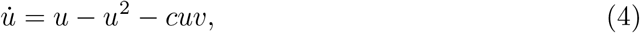

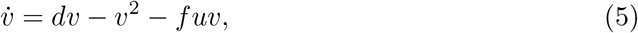

with

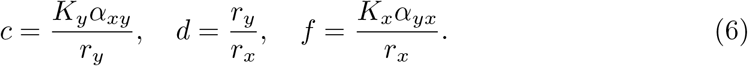

In virtue of this transformation, the equations (4) and (5) are a LV model depending upon three dimensionless parameters. This formulation is very convenient because the transformed variables are dimensionless and reduces the number of parameters, which makes easier the analysis. Furthermore, the dimensionless form of equations is extensively used in many studies, play an important role serving as parameters in differential equations, and allows describing characteristics without dimension or explicit expression unit.

The system of ordinary differential equations (4) and (5) has four equilibrium points *P*_1_(0, 0), *P*_2_(0, *d*), *P*_3_(1, 0) and 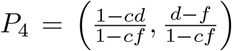. The eigenvalues of the Jacobian matrix for each of these points are given by

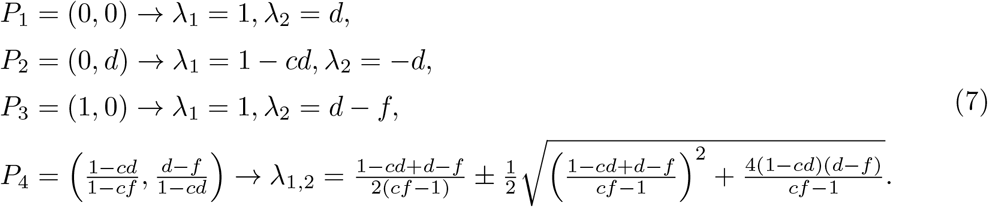

Note that in the original variables, the equilibrium points *P*_1_, *P*_2_, *P*_3_*andP*_4_ are (0, 0), (0, *K*_*x*_), (*K*_*y*_, 0) and 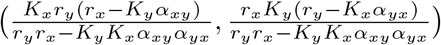, respectively.

The equilibrium point *P*_1_ is an unstable node. There exists four different regimes that are described below.

- **Regime I:** *cd >* 1 and *f* < *d* The point *P*_2_ is a stable node, and *P*_3_ is a saddle point. In the phase portrait, all the trajectories converge toward the node *P*_2_ (Figure 1a). This case may correspond to one of the types of unperturbed tumor responses that may occur naturally without the action of any external perturbation (e.g., an external anticancer therapy), named natural tumor complete remission, which may be achieved by two possible ways. First, the cancer host alone is strong enough to eliminate any unperturbed cancer formed in a long finite time. This may suppose that the psico-neuro-endocrine-metabolic-immunological system of the host (host as a whole) is strong and stable. Second, the spontaneous remission of the unperturbed tumor (tumor appears and then dissapears in a relatively short finite time). This tumor spontaneous remission may be explained because the unperturbed tumor does not find adequate biophysical conditions (metabolic, electric and energetic) in the host for its formation and growth. The duration of these two ways depends on the tumor histological variety and host type. In clinics, the cancer patient behaves like an apparently healthy individual for both ways. The absence of external perturbations, as anticancer therapies, is refered to newly diagnosed cancer patients or who do not wish to receive none anticancer therapy of their own volition. (Figure 1). The Figure 1a) is obtained using the original coordinates, taken as parameters *K*_*x*_ = *K*_*y*_ = 1 (in cells), *r*_*x*_ = 0.1 and *r*_*y*_ = 0.075 (in days^−1^), and *α*_*xy*_ = 0.1237 and *α*_*yx*_ = 0.045 (in cells^−1^× days^−1^). This gives *c* = 1.65, *d* = 0.75 and *f* = 0.45; therefore, *cd* = 1.2374 > 1 and *f* < *d* that correspond to Regime I.
- **Regime II:** *cf* < 1, *cd* < 1 and *f* < *d* The points *P*_2_ and *P*_3_ are saddle points, and *P*_4_ is a stable node. In this case, the tumor and the host coexist in equilibrium; all the trajectories converge toward the node *P*_4_ (Figure 1b). This may correspond to other two types of unperturbed tumor responses that may also occur naturally without the action of external perturbations, named natural tumor partial response or natural tumor stationary partial response. This first response type may occur when the cancer grows slowly over time (long duration response for a finite time). The second response type may occur when this coexistence remains throughout the life of the cancer patient. Therefore, the cancer may behave as a controlled disease without the need for the application of external perturbation (e.g., anticancer therapies). The last has been observed in some cancer patients in clinical oncology with long survival who are not received any type of anticancer therapy, whether oncoespecific (surgery, chemotherapy or radiotherapy) or under study/trial (e.g., immunotherapy, electrochemical therapy, irreversible electroporation) [27]. The Figure 1b) is obtained using the original coordinates, taken as parameters *K*_*x*_ = *K*_*y*_ = 1, *r*_*x*_ = 0.1, *r*_*y*_ = 0.09, *α*_*xy*_ = 0.09, and *α*_*yx*_ = 0.0788. The unities of these six parameters are previously defined. This gives *c* = 1, *d* = 0.9 and *f* = 0.788; therefore, *cd* < 1 *f* > *d* and *cf* < 1 that correspond to Regime II.
- **Regime III:** *cf >* 1, *cd >* 1 and *f* > *d* The points *P*_2_ and *P*_3_ are stable nodes, and *P*_4_ is a saddle point. The phase portrait is divided in two basin of attraction (Figure 1c); any initial state located in the upper left-hand evolves toward the node *P*_2_, whereas any trajectory in the lower right-hand side converges to node *P*_3_. In regime II, the point *P*_4_ represents a state in which the tumor and the host can coexist, but this state is unstable. This case may correspond to another type of unperturbed tumor response that may also occur naturally without the action of external perturbations, named natural tumor stable disease. The natural tumor stable disease depends on whether the endogeneous perturbation (weak or strong), either in the tumor and/or in the host, is favorable or unfavorable for the host. This coexistence is dominated by the host (node *P*_2_) when the endogeneous perturbation is favorable, whereas this coexistence is dominated by the cancer for unfavorable perturbation (node *P*_3_). In clinics, favorable endogeneous pertubation may be acceptance and positive thoughts about the disease, adequate nutrition, good performance status, among others. Unfavorable endogeneous perturbation may be stress; unpleasant news received by the cancer patient; unstabilized concomitant disease (e.g., diabetes); self-organization and dynamic transformation of the unperturbed tumor, and noise sources inherent in it and the host that occur in space-time that lead to the growth, progression, invasion and metastasis of the cancer, as well as its protection against the attack of the immune system and resistance to anticancer therapies [28, 29]. Furthermore, this self-organization leads to dynamic depletion of the host over time (most likely evolution towards node *P*_3_). The Figure 1c) is obtained using the original coordinates, taken as parameters *K*_*x*_ = *K*_*y*_ = 1, *r*_*x*_ = 0.1, *r*_*y*_ = 0.065, *α*_*xy*_ = 0.0247, and *α*_*yx*_ = 0.069. The unities of these six parameters are previously defined. This gives *c* = 1.65, *d* = 0.65 and *f* = 0.69; therefore, *cd* < 1, *f* > *d* and *cf >* 1 that correspond to Regime III.
- **Regime IV:** *cd* < 1 and *f* > *d* The points *P*_2_ and *P*_3_ are a saddle point and stable node, respectively. In clinics, it corresponds to another unperturbed tumor response that may also occur naturally without the action of external perturbation (e.g., anticancer therapies), named natural disease progression. In this case, an undesirable prognosis for the cancer patient due to his deplorable performance status (lower values of the Karnofvky index or higher value of the ECOG scale) and/or marked tumor activity (e.g., very aggressive and/or advanced-stage tumors), as shown in Figure 1d). Any initial state evolves toward node *P*_3_, i.e., the unperturbed tumor leads to energetic and metabolic depletions of the host over time that may lead to its body depauperation, which may bring about also the death of it. The Figure 1d) is obtained using the original coordinates, taken as parameters *K*_*x*_ = *K*_*y*_ = 1, *r*_*x*_ = 0.1, *r*_*y*_ = 0.066, *α*_*xy*_ = 0.066, and *α*_*yx*_ = 0.07. The unities of these six parameters are previously defined. This gives *c* = 1, *d* = 0.66 and *f* = 0.7; therefore, *cd* < 1 and *f* > *d* that correspond to Regime IV.

**Fig 1.**
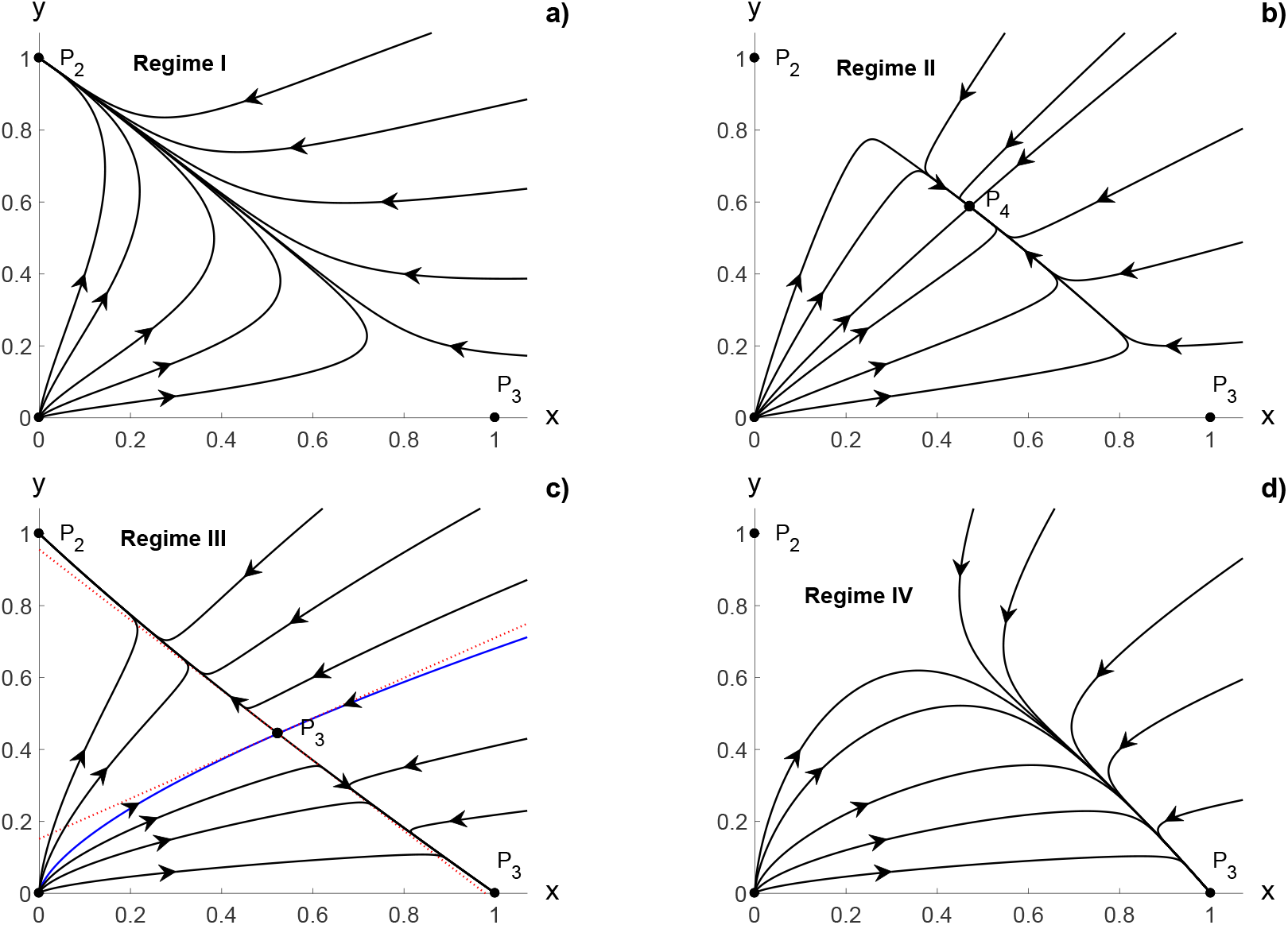
Phase portrait in the original variables: a) Regime I for *cd >* 1 and *f* < *d*. b) Regime II for *cf* < 1, *cd* < 1 and *f* < *d*. c) Regime III for *cf >* 1, *cd >* 1 and *f* > *d*. d) Regime IV for *cd* < 1 and *f* > *d*.

From the analysis performed in this Section 2, it is clear that there is not oscillating solution. The existence of periodic solutions is treated in the Section 3 of this study. Strictly speaking, the assumption of constant parameters in our model is only an approximation valid inside certain time scales. Many processes in the host and its environment are cyclic (e.g., concentration of nutrients and oxygen, elimination of metabolic byproducts). In addition, cancer cells may develop control mechanism to modify the environment [4–8], which yield to a modification of the parameters. In spite of this limitation, the model describes adequately the different dynamic regimes of tumor evolution.

An important merit of this model is that there is not unlimited growth, i.e., all the trajectories in the phase portrait are bounded [9], in agreement with the experiment [27, 28, 30–32] an other theoretical studies [33]. Although the transformation performed in this study greatly simplifies the analysis of the system of equations (4) and (5), with respect to those of equations (1) and (2), another merit of the transformed model is that this transformation does not change the dynamic regimes nor nature of tumor-host interaction. Furthermore, although the parameters *c, d* and *f* are dimensionless in equations (4) and (5), these three parameters depends on all the biologically meaningful original parameters (*r*_*x*_, *K*_*x*_, *α*_*xy*_, *r*_*y*_, *K*_*y*_ and *α*_*yx*_ in equations (1) and (2)). Therefore, the parameters *c, d* and *f* sense the changes in six original parameters.

The clinical interpretation of these four dynamic regimes and their natural tumor responses associated depend on degree of immunocompetence (immunocompetent or immunodeficient) of the host, the tumor histological variety, and endogenous perturbations inherent in the cancer and host. In this case, favorable endogenous perturbations may behave as natural anticancer therapies directed at the host, so as to induce metabolic, energetic and physiological conditions unfavorable to the cancer. This may lead to its natural tumor complete remission or natural tumor stationary partial response. If we would have a good understanding of these aspects, the cancer cure would be physiological (withouth the need fon anticancer therapies), idea that should not be underestimate. This would imply that scientific thinking should be directed to the cancer patient as a whole and not to the cancer itself.

### About the existence of periodic orbit

The criterion of Dulac [34, 35] states that if the function *R*(*x, y*) is defined on a simply connected region *S* ⊆ ℝ^2^ and the expression 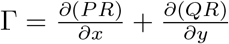 is not identical to zero and does not change sign, then the system of equations

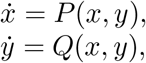

has no closed orbits lying entirely in *S*. By taking the function of Dulac *R* = *u*^*m*^*v*^*n*^ [33], *m* = (*cf* + *f* − 2)*/*(1 − *cf*) and *n* = (*cf* + *c* − 2)*/*(1 − *cf*), we get

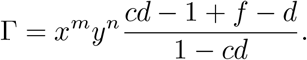

If *cd* − 1 ≠ *d* − *f*, then Γ = 0 only when *x* = 0 or *y* = 0. For the regimes II and III, we have that (*cd* − 1 + *f* − *d*)*/*(1 − *cf*) < 0, then the function Γ do not change its sign inside of any of the quadrants (|*u*|, |*v*| > 0).

For the regimes I and IV, the condition Γ = 0 is fulfilled when *cd* − 1 = *d* − *f*. In such case, the fixed point *P*_4_ has purely imaginary eigenvalues when *cf* > 1, given by

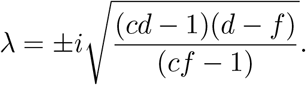

Since there are not periodic solution in the first quadrant and this system is constrained to nonnegative values of the variables (*u, v*) ≥ 0, there are not periodic solutions for this model.

### Cancer and host cell populations under an external perturbation

Although the four regimens for dynamic unperturbed tumors reported in this Section 1 have been reported previously by several authors [7–9], we do not know how these regimes change when the host, not the tumor, is perturbed with external perturbation (e.g., anticancer therapies) and their possible correspondence with results reported in clinical and experimental oncology, as in this Section 4. Furthermore, the clinical interpretation of the four dynamic regimes and types of natural tumor responses should not be confused with those reported when external perturbations, as anticancer therapies are used.

It is well known that the application of oncospecific anticancer therapies (e.g., chemotherapy and radiotherapy) damages both the cancer and the host (specifically the healthy tissue surrounding the tumor) [9, 36, 37]. In this case, equations (1) and (2) are modified by these therapies, given by

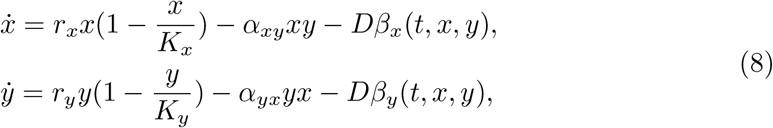

where *D* is the relative dose (dimensionless parameter), which is defined as the ratio between the dose applied to the host with cancer and the maximum permissible dose that the host can receive for the anticancer therapy used. We assume that the effect that this therapy produces in both systems is proportional to *D*, which produces destroys *Dβ*_*x*_(*t, x, y*) cancer cells and *Dβ*_*y*_(*t, x, y*) normal cells per unit of time. The *β*_*x*_(*t, x, y*) and *β*_*y*_(*t, x, y*) are given in cells/days.

The transformed system (4), (5) becomes in

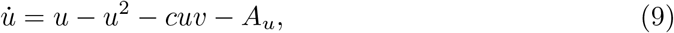

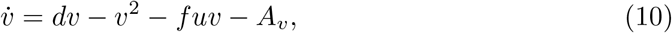

with

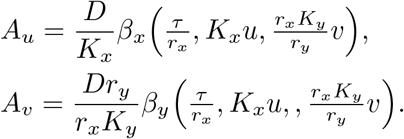

In the case that the effect of the therapy on the tumor cells does not depend on the healthy cells, that is *β*_*x*_ = *β*_*x*_(*t, x*), from equation (9), we can write

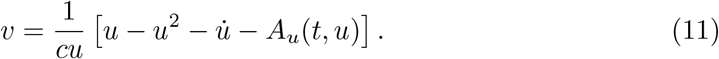

By inserting (11) into (10), it is obtained that

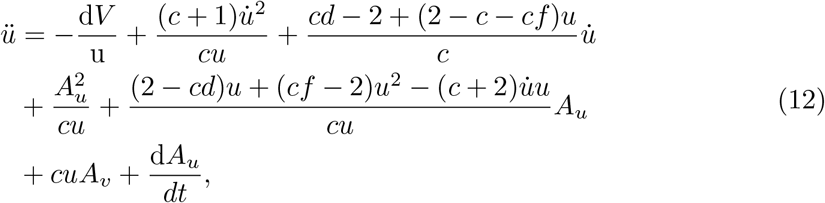

where

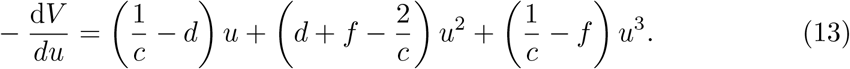

Equations (12) and (13) may be interpreted as the motion of a particle in a viscous medium under the action of a force field [24, 25]. In the case that *A*_*u*_(*t, u*) = *A*_*v*_(*t, v*) = 0, the equilibrium points are determined by the potential *V* (*u*). The potential function has three equilibrium points that are determined from the equation *dV* (*u*\*)*/du* = 0: *u*_1_* = 0, *u*_2_* = 1, *u*_3_* = (1 − *cd*)*/*(1 − *cf*). The condition of stability is given by

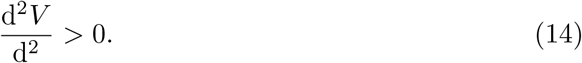

By evaluating (14), we find that *u*_1_* = 0 is stable for *cd* > 1; *u*_2_* = 1 is stable for *f* > *d*; *u*_3_* = (1 − *cd*)*/*(1− *cf*) is stable for *f* < *d*. Note that *V* (*u*) is a polinomial of fourth degree that has three real exrema., two minima and one maximum if *cf* > 1 or two maxima and one minimum if *cf* < 1. There are four possible distinctive cases for the potential depending on the values of the parameters *c, d* and *f* (Figure 2).

**Fig 2.**
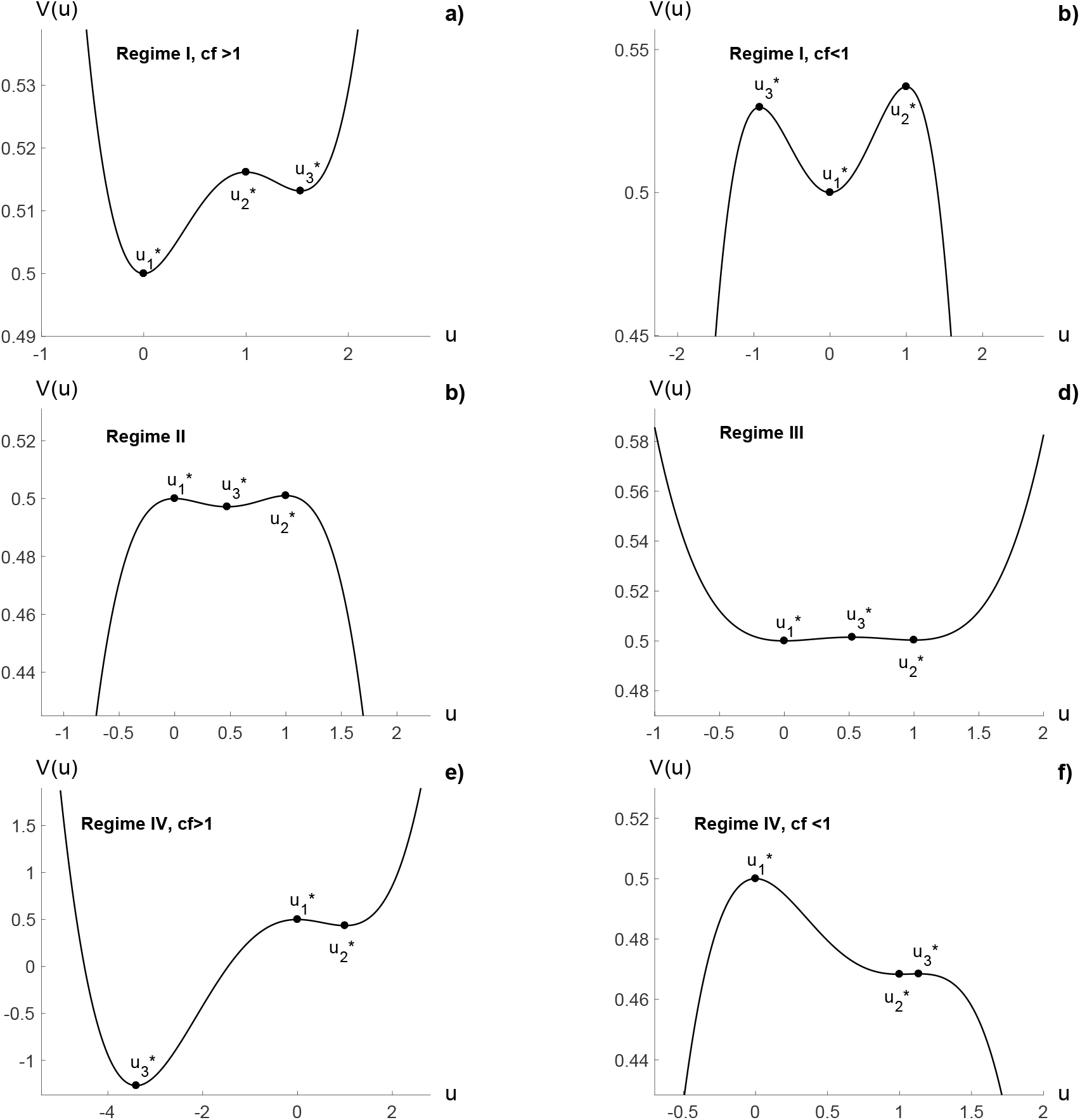
The potential function *V* (*u*): a) *cd* > 1, *f* < *d* and *cf* < 1 b)*cd* > 1, *f* > *d* and *cf* > 1 c) *cd* < 1, and *f* > *d*, d) *cd* > 1 and *f* > *d*, e) *cd* < 1, *f* > *d* and *cf* > 1, f) *cd* < 1, *f* > *d* and *cf* < 1. The value of the parameters are the same as those used in Figure 1 for b), c), d) and f) subfigures. Subfigure a) was obtained with *c* = 1.65, *d* = 0.75 and *f* = 0.7. Subfigure e) was obtained with *c* = 1, *d* = 0.66, and *f* = 1.1

- **Case 1**.: *cd* > 1 and *f* < *d*
  - If *cf* > 1, *u*_1_* and *u*_3_* are minima, *u*_2_* is a maximum (Figure 2a).
  - If *cf* < 1, *u*_2_* is a minimum, *u*_2_* and *u*_3_* are maxima, (Figure 2b).
- **Case 2.;** *cd* < 1 and *f* < *d* (*cf* < 1)
  - *u*_3_* is a minimum, *u*_1_* and *u*_2_* are maxima (Figure 2c).
- **Case 3:** *cd* > 1 and *f* > *d* (*cf* > 1)
  - *u*_1_* and *u*_2_* are minima, *u*_3_* is a maximum (Figure 2d).
- **Case 4:** *cd* < 1 and *f* < *d*
  - If *cf* < 1, *u*_2_* is a minimum, *u*_1_* and *u*_3_* are maxima, (Figure 2e).
  - If *cf* > 1, *u*_2_* and *u*_3_* are minima, *u*_1_* is a maximum (Figure 2f).

Notice that the Cases 1, 2, 3 and 4 correspond to the Regimes I, II, III and IV respectively. Unlike the four regimens for unperturbed tumors (see Section 1 of this study), the four regimens shown in this Section are for tumors under an external perturbation. These new regimes depend on the performance status of the cancer patient, tumor histological variety and doses *D* of therapy applied to both the cancer and host.

The LV model and the particle-like equation (12) do not have periodic solutions; nevertheless, the particle may oscillate around the stable equilibrium positions under a periodic driving force. It is possible to define a characteristic or proper frequency around the minima of *V* (*u*) as

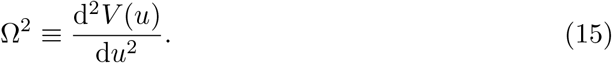

In the case of Regime IV, we have that Ω^2^ = *f* − *d* for *u*\* = 1.

The behavior of periodic forced nonlinear systems is well known for low dimensional systems [38, 39]. The motion is synchronized to the external force for frequencies *ω* of the external force close to Ω [38]. In general, the amplitude of the oscillation decreases as the difference *ξ* =| Ω− *ω*| increases and the synchronization is destroyed for higher values of *ξ*, resulting in a more complex motion that in some cases may be chaotic.

### Cancer and host cell populations under a continuous external perturbation

The continuous application of external perturbation to the cancer and host implies that chemotherapy and radiotherapy cannot be applied continuously over time because of their marked adverse events [9, 36, 37], not so for immunotherapy that is applied to the cancer patient throughout life [40–42]. Nevertheless, several studies report immune-related adverse events that may harm the host. The adverse events induced by immunotherapy in the host are less than those induced by chemotherapy or radiotherapy [43–45].

It is important to note that destruction of healthy host cells is negligible compared to that induced in the tumor, so *β*_*x*_(*t, x, y*) >> *β*_*y*_(*t, x, y*). Otherwise, external perturbation cannot be used because it would induce important adverse events in the host, including death. It is well know that safety and adverse events, in addition to efficacy/effectiveness, are very important therapy aspects aimed at any type of disease must fulfill simultaneously, specifically anticancer therapies.

### Cancer and host cell populations affected proportionally under a continuous external perturbation

We assume that the external perturbation induces destruction of cancer cells and host cells at constant rate (same amount of cells per unit of time), as long as *β*_*x*_(*t, x, y*) >> *β*_*y*_(*t, x, y*). In this case, *β*_*x*_(*t, x, y*) and *β*_*y*_(*t, x, y*) do not depend on time *t*, that is, *β*_*x*_(*t, x, y*) = *β*_*x*_*x* and *β*_*y*_(*t, x, y*) = *β*_*y*_*y*. In such case, the LV equations (12) and (13) reduce to

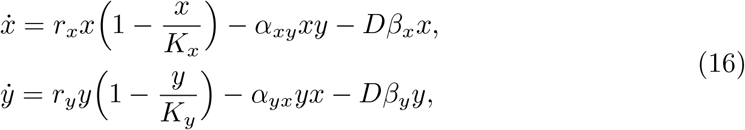

This is again a LV system with growth factors *r*_*x*_ − *Dβ*_*x*_, *r*_*y*_ − *Dβ*_*y*_ and carrying capacities *K*_*x*_(*r*_*x*_ − *Dβ*_*x*_)*/r*_*x*_, *K*_*y*_(*r*_*y*_ − *Dβ*_*y*_)*/r*_*y*_.

If *r*_*x*_ −*Dβ*_*x*_ ≤ 0, then 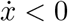. This implies that *x* is stricttly decreasing and *x* → 0 with time. The external perturbation destroys the tumor completely. Therefore, the tumor will be destroyed for highly enough dose *D*. Unfortunately, high dose may destroy also many healthy cells. Consequently, adverse events are induced in the host that may even be severe and subsequently lead to its death.

If *r*_*y*_ − *Dβ*_*x*_ ≤ 0, then 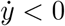 and *y* → 0 with time. The external perturbation destroys the healthy cells completely. This will not be a problem if the destroyed cells restrict to a limited, non essential, area of the body (e.g., healthy tissue surrounding the tumor), as induced by electrochemical ablation [27, 28, 30–32].

Let us assume that *D* < min(*r*_*x*_*/β*_*x*_, *r*_*y*_*/β*_*y*_). Since *r*_*x*_ − *Dβ*_*x*_ > 0 and *r*_*y*_ − *Dβ*_*x*_ > 0, the equations can be transformed to a system (4), (5) with coefficients

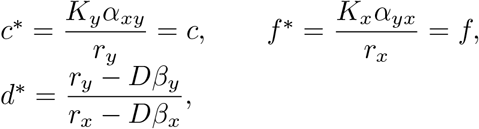

and with the transformed variables defined by

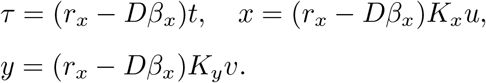

Let us analyze the condicions in which the external perturbation moves the system to a Regime that takes the cancer patient to a cured state (cancer patient behaves like an apparently healthy individual). We assume that the system is in Regime IV (*cd* < 1 and *d* < *f*) or in Regime III with inadequate initial conditions (*cf* > 1, *cd* > 1 and *f* > *d*).

We wish to move the system to Regime I, Regime II, or even to Regime III for the case of suitable initial conditions (the cancer patient state under external perturbation application). Note that a variation in the dose *D* only affects the coefficient *d*\*. Then, the external perturbation is effective only if *d*\* is an increasing function of *D* and the first derivative of *d*\* with respect to *D* must be positive (*r*_*y*_*β*_*x*_ − *r*_*x*_*β*_*y*_ > 0 or *β*_*x*_*/β*_*y*_ > *r*_*x*_*/r*_*y*_).

As usually *r*_*x*_ ≫ *r*_*y*_, *β*_*x*_ ≪ *β*_*y*_. This means in the therapeutic order that the effect of the external perturbation on tumor cells must be much larger than the effect on healthy cells in a proportion at least as the proportion between the growth ratio of the tumor cells over the growth ratio of the healthy cells.

- Cancer patient in Regime IV (*cd* < 1 and *f* > *d*) Condition *cd* < 1 is equivalent to *r*_*x*_ > *K*_*y*_*α*_*xy*_ and condition *f* > *d* is equivalent to *r*_*y*_ < *K*_*x*_*α*_*yx*_. This implies that

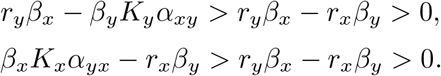 In this situation,

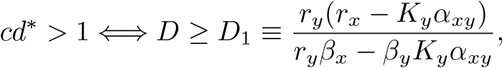

and

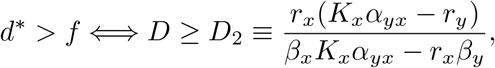

then,
  - If *D* > max(*r*_*x*_*/β*_*x*_, *r*_*y*_*/β*_*y*_), tumor cells and healthy cells go to zero.
  - If *D* > max(*D*_1_, *D*_2_), the system goes to Regime I.
  - If *cf* < 1 and *D*_1_ > *D > D*_2_, the system goes to Regime II.
  - If *cf* > 1 and *D*_2_ > *D > D*_1_, the system goes to Regime III.
  - If *D* < min(*D*_1_, *D*2), the system stays in Regime IV.
  In all the cases, if the external perturbation is withdrawn, the system returns to Regime IV, unless the tumor has been completely destroyed. Note that in the mathematical model this is achieved only when *t* → ∞. Nevertheless, in practice this may be accomplished if the variable *x* is smaller than a small quantity.
- Cancer patient in Regime III (*cf* > 1, *cd* > 1 and *f* > *d*) Since *cd* > 1 and *d*\* is an increasing function of *D*, the condition *cd*\* > 1 is satisfied for any *D* ≥ 0. The condition *f* > *d* is equivalent to *r*_*y*_ < *K*_*x*_*α*_*yx*_ and this implies that

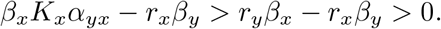 In this case,
  - If *D* > max(*r*_*x*_*/β*_*x*_, *r*_*y*_*/β*_*y*_), tumor cells and healthy cells go to zero.
  - If *D* > *D*_2_, the system goes to Regime I.
  - If *D* > *D*_2_, the system stays in Regime III.
  The system returns to Regime III when the application of external perturbation is finished, unless the values of *x, y* at this moment are in the region of the plane (*x, y*) in which the solution of (4),(5) tends to the equilibium point *P*_2_, that is, if *x* is small enough and *y* is large enough.
- Cancer patient in Regime II (*cf* < 1, *cd* < 1 and *f* < *d*) Since *d* > *f*, the condition *cd*\* > 1 is satisfied for any *D* ≥ 0. Condition *cd* < 1 is equivalent to *r*_*x*_ > *K*_*y*_*α*_*xy*_ and this implies that

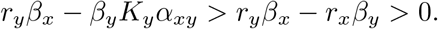 In this case
  - If *D* > *D*_1_, the system goes to Regime I.
  - If *D* > *D*_1_, the system stays in Regime II.
  The system returns to Regime II when the application of the external perturbation is finished, unless the values of *x* at this moment are small enough to be considered zero, that is, the tumor has been destroyed.

Nowadays, anticancer therapeutic strategies has been directed primarily at cancer with host damages, which may be mild, moderate or severe depending on the type of therapy applied [9, 36, 37, 40–46]. Nevertheless, we are not aware that anticancer therapeutic strategies are directed solely at the host in order to create unfavorable conditions in it that lead to complete remission or stationary partial remission of the cancer without adverse events in the host or if these are induced in the host, they are minimal (the host recovers in the shortest possible time). This may be achieved with a constant (see Subsection 4.3) or periodic (see Subsection 4.4) external perturbation.

### Host cells population under an external perturbation applied continuosly

If we assume that the external perturbation destroys a constant number of tumor cells and host cells per unit of time, the system (8), can be rewritten as

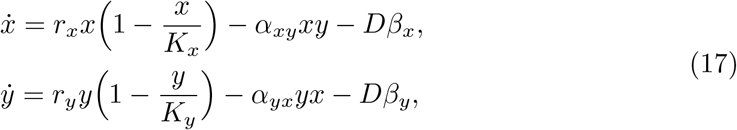

As the intention of this subsection is to analyze the system when the host is only perturbed with a constant external perturbation, the case considered is the following: *β*_*x*_ = 0 and *β*_*y*_ < 0. The effect of the constant term is studied by Kim [23] to explain the effect of harvesting on the dynamic of two competing species. For small values of *A*, the system only undergoes a slight distortion in the phase portrait with a minuscule shift in the equilibrium points. Using the change of variable (3) and the change of parameters (6), the equations (17) transform to

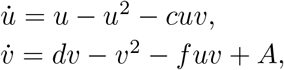

with 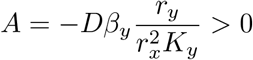.

As in the case of previous equations, if the initial conditions *u*(0), *v*(0) are non negative, the solution stays in the positive quadrant, as expected in clinics since no negative populations can happen. Then, only the positive quadrant is considered.

For this system (equations (25) and (26)), we find the following equilibrium points

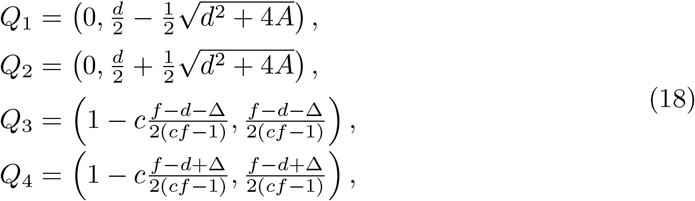

where 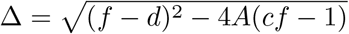.

For small values of *A*, the fix points are moved slightly with respect to those corresponding to the unperturbed tumor case (Figure 1), as shown in Figure 3. An important fact is that the point (*x*, 0) is never an equilibrium point. The population of healthy cells cannot vanish. The Regimes IV* is equivalent to Regime II*, in which both cancer and healthy cells coexist in the equilibrium point. The size of the equilibrium populations depend on the initial state (Regime II or Regime IV) and on the applied therapy dose.

**Fig 3.**
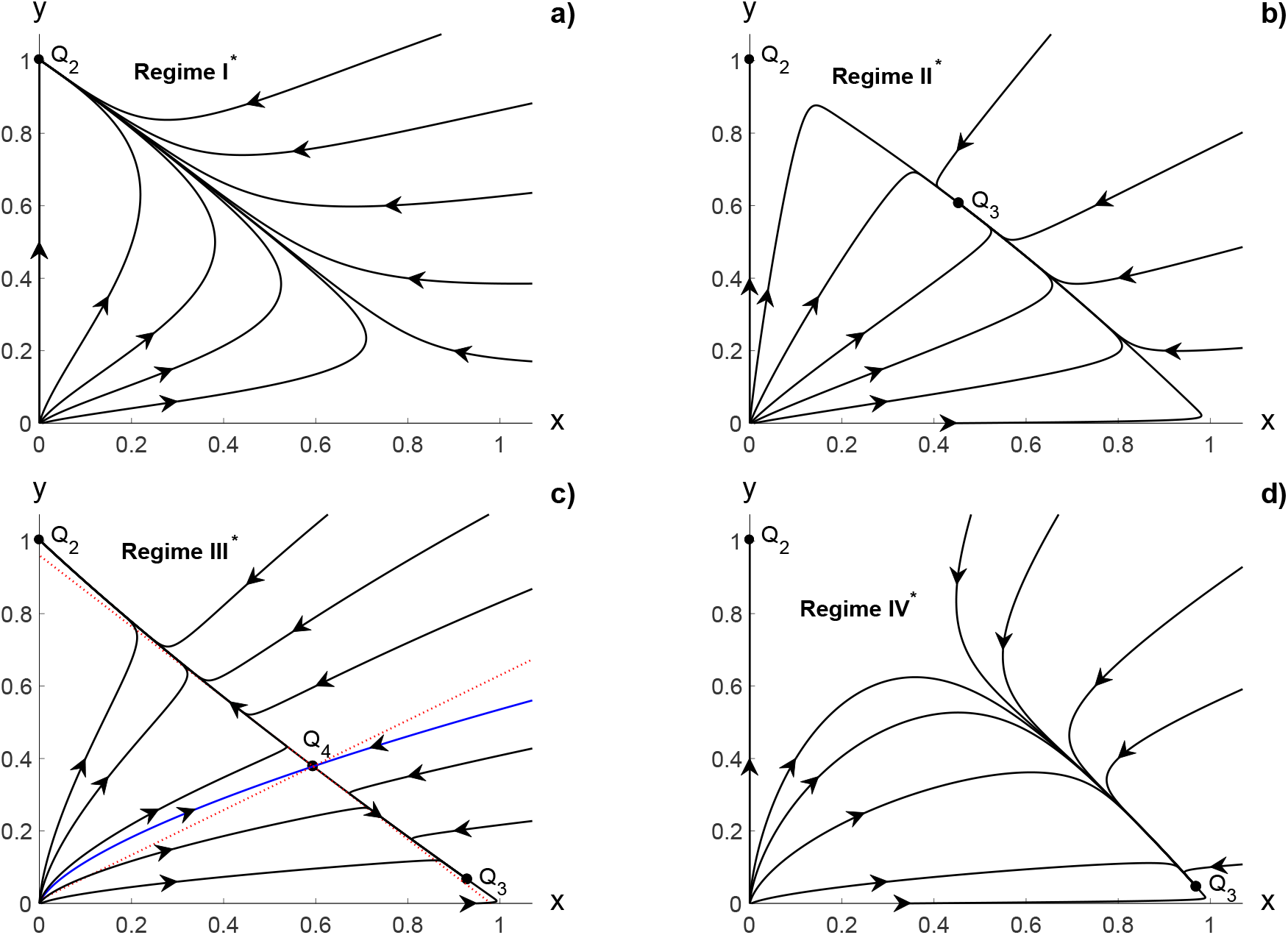
Effect of external stimulus with low continuous dose, in the original coordinates (17): a) on Regime I, b) on Regime II, c) on Regime III, d) on Regime IV.

The dislocation can be considerable and bifurcations can occur for higher values of *A*, as we will expose below.

The point *Q*_1_ has eigenvalues 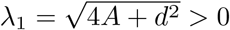 and 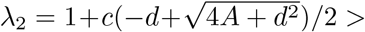 0 and is always unstable. The point *Q*_2_ has eigenvalues 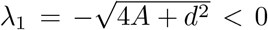 and 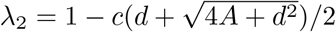 and is stable for *A* > *A*_1_ with

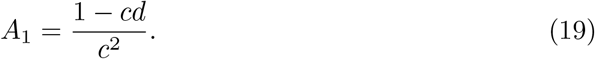

In terms of the original parameters, it is stable when

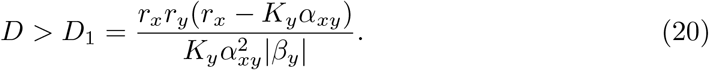

In order for the fixed points *Q*_3_ and *Q*_4_ to exist, the expression inside square root of Δ must be nonnegative. Since *A* > 0, this is always true if *cf* ≤ 1, as happens in Regime II and also if *cf* > 1 and *A* ≤ *A*_2_ with

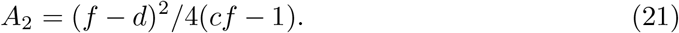

In terms of the original parameters, this happens when

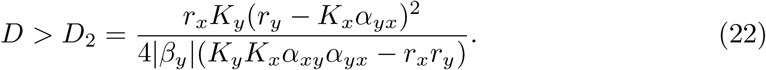

Moreover, if (*f* − *d*)(*cf*− 1) < 0, since Δ ≤ |*f*− *d*|, the points *Q*_3_ and *Q*_4_ have the second component negative and have no interest from the clinical point of view. The equilibrium points *Q*_3_ and *Q*_4_ make sense only when *A* ≤ *A*_1_, *cf* > 1 and *d* > *f* or well when *cf* < 1 and *d* > *f*. As *A* → *A*_1_, the points *Q*_3_ and *Q*_4_ approach until they collide when *A* = *A*_1_ creating a new equilibrium point, which has the eigenvalues *λ*_1_ = 0 and 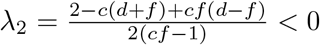. The point 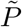 disappears for higher values of *A*.

- Cancer patient in Regime IV (*cd* < 1 and *f* > *d*) In Regime IV and for *cf* > 1, a bifurcation occurs when *A* = *A*_2_ (*D* = *D*_2_), for which *Q*_3_ and *Q*_4_ collide. If *A* > *A*_2_ (*D* > *D*_2_) there are only two equilibrium points: *Q*_1_ is an unstable node whereas *Q*_2_ is stable (it can be verified that *A*_1_ < *A*_2_ in this case) and the patient goes to a Regime I*. If *A*_1_ < *A < A*_2_ (*D*_1_ < *D < D*_2_) there are two possibilities: If *c*(*f* −*d*) > 2(*cf*− 1), the first component of *Q*_4_ and *Q*_3_ are negative, and the patient goes to a Regime I*. If *c*(*f* −*d*) < 2(*cf* −1), *Q*_3_ and *Q*_4_ are in the positive quadrant, and the patient goes to Regime III*. Finally, if *A* < *A*_1_ (*Q*_2_ and *Q*_4_ collide at *A* = *A*_1_), the point *Q*_2_ is not stable but *Q*_3_ is, and the patient goes to Regime II*. When *cf* < 1 on regime IV, the second component of point *Q*_4_ is negative and this point has no interest from the biological point of view. Moreover, a bifurcation occurs when *Q*_2_ and *Q*_3_ collide, which happens when *A* = *A*_1_. For *A* > *A*_1_ (*D* > *D*_1_), the first component of *Q*_3_ is negative and there are only two equilibrium points: *Q*_1_ is an unstable node whereas *Q*_2_ is stable and the patient goes to Regime I*. If *A* > *A*_1_, *Q*_2_ is not stable but *Q*_3_ is stable. The patient goes to Regime II*. In conclusion,
  - If *D* > max(*D*_1_, *D*_2_), the system goes to Regime I*.
  - If *cf* < 1 and *D* > *D*_1_ (*D*_2_ < 0), the system goes to Regime II*.
  - If *cf* > 1, *c*(*f* − *d*) > 2(*cf* − 1) and *D* > *D*_1_, the system goes to Regime I*.
  - If *cf* > 1, *c*(*f* −*d*) < 2(*cf*− 1) and *D*_2_ > *D > D*_1_, the system goes to Regime III*.
  - If *D* > *D*_1_, the system goes to Regime II*.
- Cancer patient in Regime III (*cd* > 1, *cf* > 1 and *f* > *d*) In this regime, *A*_1_ < 0, therefore, *Q*_2_ is always stable. If *A* < *A*_2_, *Q*_3_ and *Q*_4_ are in the positive quadrant, *Q*_4_ is not stable and *Q*_3_ is stable. The patient goes to Regime III*. If *A* > *A*_2_, the patient goes to Regime I*. In conclusion,
  - If *D* > *D*_2_, the system goes to Regime I*.
  - If *D* > *D*_2_, the system goes to Regime III*.
- Cancer patient in Regime II (*cd* < 1, *cf* < 1 and *f* < *d*) In this regime, *Q*_4_ is never in the positive quadrant. If *A* < *A*_1_, *Q*_3_ is stable but *Q*_2_ is not, and the patient goes to Regime II*. If *A* > *A*_1_, *Q*_2_ is stable and *Q*_3_ has the first component negative. The patient goes to Regime I*. In conclusion,
  - If *D* > *D*_1_, the system goes to Regime I*.
  - If *D* > *D*_1_, the system goes to Regime II*.

The results of this Subsection demonstrate that it is possible to obtain complete tumor remission for sufficiently high doses of a constant external perturbation. This is impressive because a very aggressive tumor may be destroyed not by direct attacking it but enhancing the host (e.g., stimulating the immune system) using high constant doses of an external perturbation. Unfortunately, such powerful stimulating anticancer therapies are not available at present due to the possible induction of adverse events in the host. This is true for currently existing anticancer therapies.

Although a constant external perturbation over a host may induce complete tumor remission, its exposure time may be short or long depending on whether the adverse events in the host are marked (irreversible tissue damage or death) or not. If adverse events are marked, the exposure time of this constant external perturbation is of short duration; otherwise, it is of long duration.

In this study, high dose is referred to that applied to the host in such a away as to sufficiently potentiate the immune system and/or create unfavorable conditions in the host in order to induce complete remission or stationary partial response of the tumor through the apoptosis mechanism without provoking severe adverse events in the whole organism. This may suggest the tumor sensitivity to the external perturbation applied to the host must be much greater than that of the rest of the organism. In other words, the tissue damage induced in the tumor is amplified several orders of magnitude compared to that of the rest of the organism. This would represent a new concept for cancer therapy, unprecedented in the literature.

In the therapeutic order, the external constant perturbation may confirm that the target of anticancer therapies should be the different types of cells that compose the tumor stroma, rather than the cancer cells, for the long-term control of this disease [47–49]. In principle, this external perturbation type may be a biological or physical therapy, which may be combined. Among the biological therapies may be mentioned anti-VEGF therapy (antiangiogenic therapy) [50–53], immunotherapy using immune checkpoint inhibitors (anti-chekpoint therapies) [54–56], combination of antibody therapies [57, 58]. Nevertheless, the use of these anticancer therapies may induce severe adverse events [59–61]. On the other hand, constant electric stimulation therapies applied directly to the host for restoration of the tumor electrical microenvironment [62–66]. This may lead to modifications of the tumor stroma, either by repolarization of its components, reestablishment of the pH of the environment, inhibition of blood vessels, among others. Electrical therapies should not be confused with those applied directly to the tumor (e.g., electrochemical ablation) [27, 28, 30–32].

Consequently, these tumor stroma-targeted anticancer therapies need to be redesigned or new therapeutic strategies proposed. A more reasonable therapeutic strategy is the application of an external stimulus applied periodically to host cells population. This may be modelled by taking the applied dose as a (periodic) function of time, or equivalently taking *β*_*x*_ and *β*_*y*_ periodical functions of time.

### Host cells population under an external perturbation applied periodically

As a first approximation we will assume that *β*_*x*_(*t, x, y*) = *D*(1 − cos(*ωt*)*β*_*x*_ and *β*_*y*_(*t, x, y*) = *D*(1 − cos(*ωt*)*β*_*y*_. In such a case, the LV equations reduce to

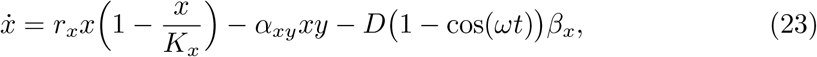

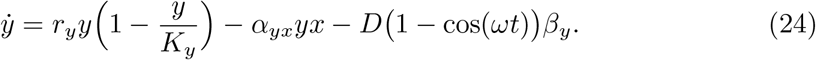

Moreover, we will consider here the case where the external stimulus applied periodically affects only the host cells, with *β*_*y*_ < 0. Then the system (23),(24), reduces to

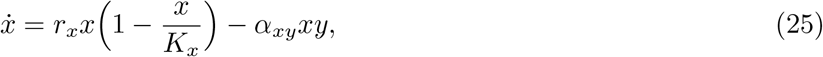

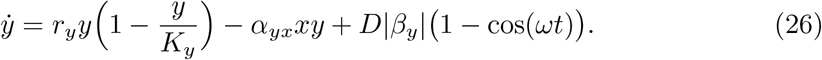

An analytical study is not easy, so simulations are performed by numerically integrating these systems of differential equations for regime IV. A limit cycle is obseved around the position where *Q*_3_ was located (see Figure 4a)). The oscillations of periodically forced systems synchronize to the driving term and the amplitude of oscillations decreases with the increase of the frequency *ω* (these occurs in a certain range of *ω* around the characteristic frequency Ω (15) of the unperturbed system) [33, 34], as it is shown in Figures 4b) and 4d). For higher values of *ω* the limit cycle becomes an attractor (see Figure 4c)).

**Fig 4.**
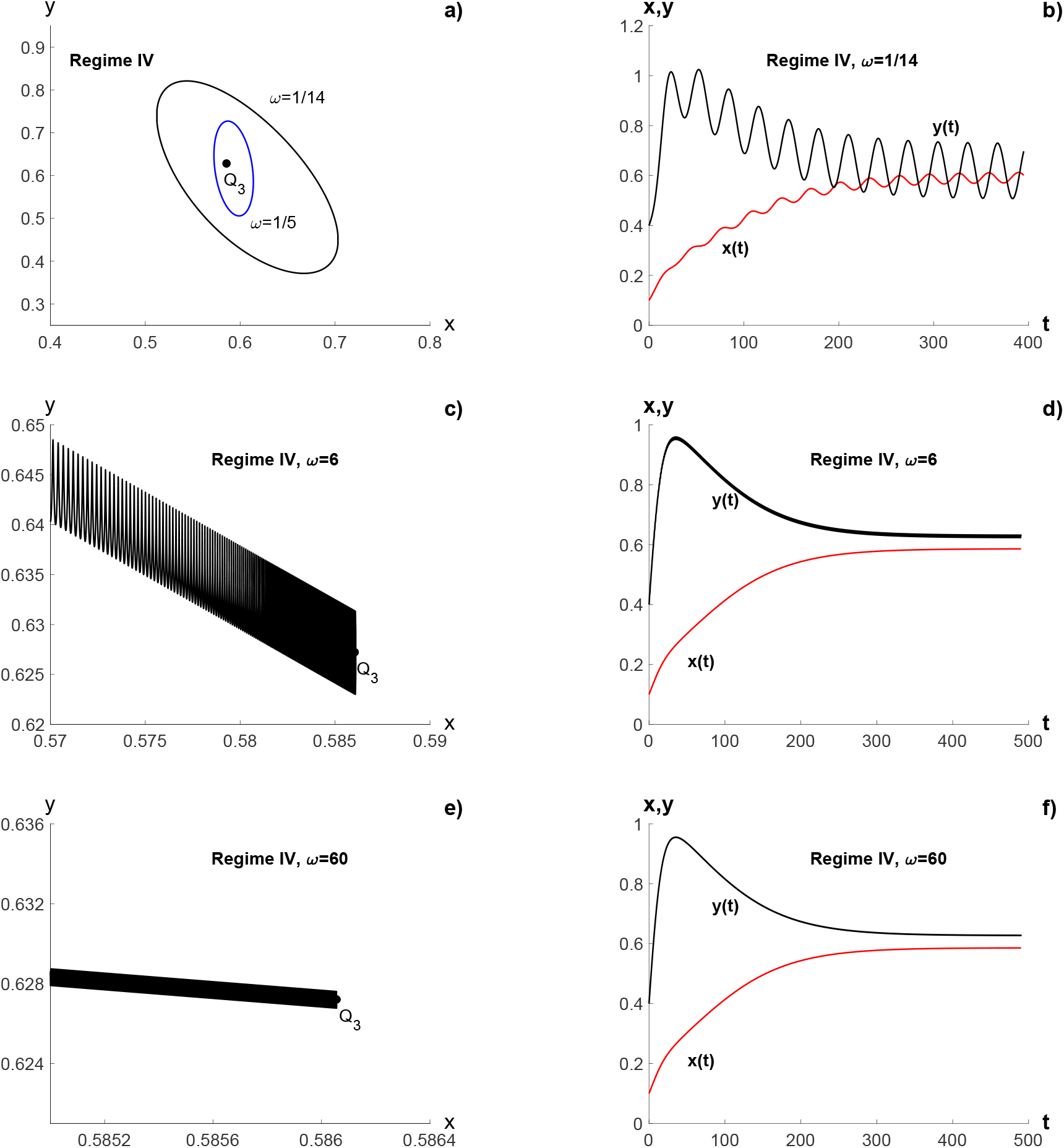
System on regime IV under a periodic perturbation, in original coordinates: a) limit cycle for *ω* ∼ Ω; b) Solutions for *ω* = 1*/*14; c) limit cycle for *ω* = 6; d) Solutions for *ω* = 6; e) non-chaotic attractor for *ω* ≫ Ω; f) solutions for *ω* = 60 The parameters values are *c* = 1, *d* = 0.66, *f* = 1.1, *A*_1_ = 0, which gives Ω = 0.2, and *A* = 0.175.

The subfigures in Figure 4 are obtained using the original coordinates, taken as parameters *K*_*x*_ = *K*_*y*_ = 1, *r*_*x*_ = 0.1, *r*_*y*_ = 0.066, *α*_*xy*_ = 0.066, *α*_*yx*_ = 0.11., *β*_*y*_ = 1*/*40 and *D* = − 1. This gives *c* = 1, *d* = 0.66 and *f* = 1.1; therefore, *cd* < 1 and *f* > *d* that correspond to Regime IV.

We calculate the maximum Lyapunov exponents to characterize this attractor [35], defined as

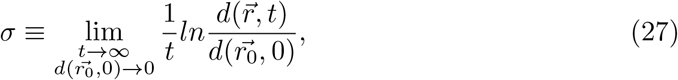

where 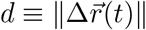 is the distance between two initially close trajectories, that is, the trajectories where at 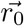 and 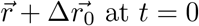. The evolution of distance between both trajectories is calculated from the linearized system

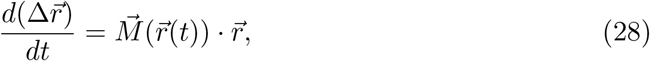

where 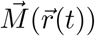 is the Jacobi matrix of the system (25)–(26) evaluated along a fix trajectory 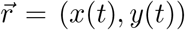. To avoid numerical overflow of the distance 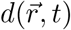, it is chosen a small fixed time interval *τ*, and *d* was renormalized to *d*_0_ every *τ*. Thus, we iteratively compute the values

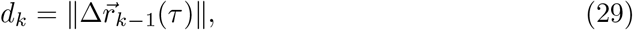

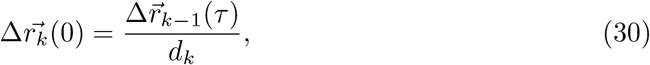

where 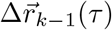 is obtained by integrating (28), with the initial value 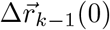. Defining the quantity

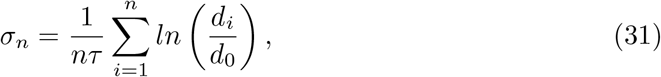

it can be shown that for a *τ* not too large lim_*n*→∞_ *σ*_*n*_ = *σ* and is independent of *τ*.

Figure 5a) shows *σ*_*n*_ versus *n* for *ω* = 0.5, where convergence can be seen. The dependence of *σ* on *ω* is shown in Figure 5b). The maximum Lyapunov exponent remains negative for high frequencies, such as *ω* = 300.

**Fig 5.**
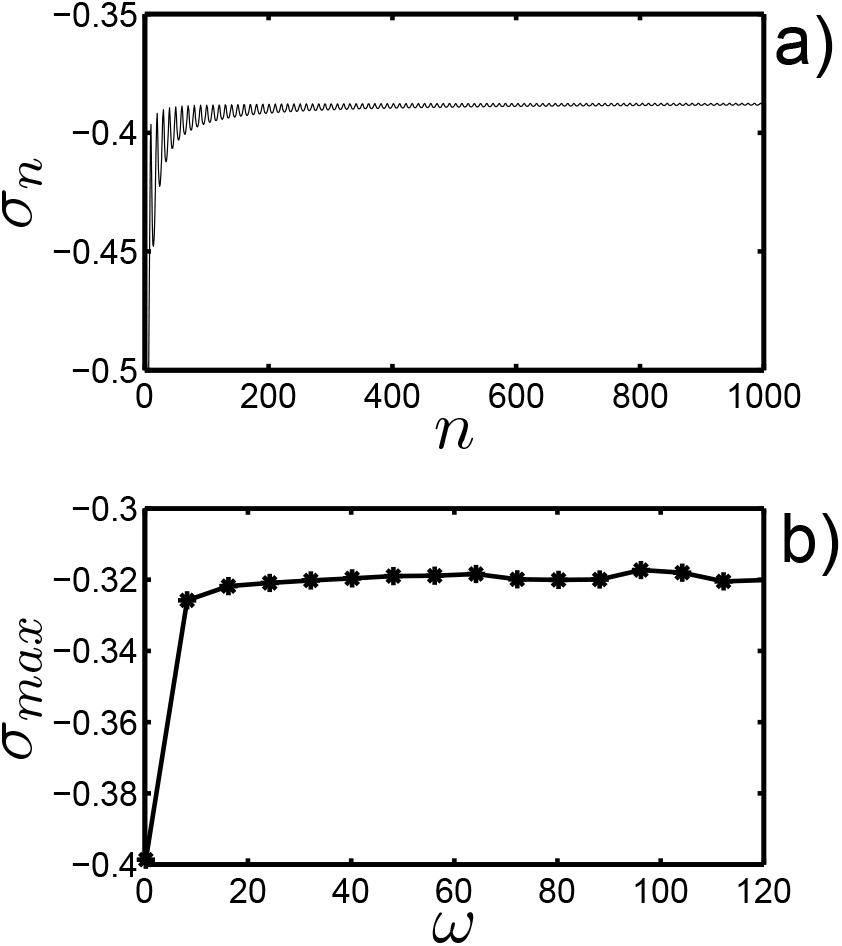
Lyapunov exponents: a) *σ*_*n*_ vs. *n*; and b) dependence of maximum Lyapunov exponent with the frequency. The parameters are the same of Figure 4.

In the therapeutic order, the external periodic perturbation may be of a biological or physical nature that may be combined as well. There exist moderated stimulating biological anticancer therapies whose action are not constant, but may be applied to the patient periodically, such as the use of cytokines and vaccines to enhance the response of the immune system [38, 67–71]. Furthermore, physical therapies may be an electromagnetic field at certain frequencies (alternating electric fields and/or a periodically applied pulsed electromagnetic field) that may harmonize the host (restoration of injury and disorders caused by the tumor in its vecinity), modulate the tumor stroma, stimulate the immune system response and inhibits tumor angiogenesis in order to induce the complete remission or stationary partial response (limit cycle: cancer as a controlled chronic disease). For this, the ideas reported in several studies must be taken into account [72–78].

## Summing up and outlook

When the oncoespecific anticancer therapies, immunotherapy and physical therapies are used, the complete remission of highly aggressive tumors is observed less frequently. This also occur for advanced tumor stages. This tumor response type does not always mean the cancer is completely cured, aspect that may be explained because some cancer cells may remain in body for many years after application of these therapies. Furthermore, these therapies reduce the size of these tumors, but they may regrow at a later time after the application of these external perturbations directed directly to the tumor. In the worst scenario, these therapeutic strategies may accelerate the death of the patient.

The LV model is used to test two different therapeutic strategies. First, we study the action of a constant external perturbation to healthy cells of the host. In principle, it should be possible to eradicate the malignant cells choosing the appropriate doses and exposure time. Unfortunately, such external perturbations are not available in clinical or experimental oncology at present. This may be due to their doses and exposure time are limited by the tolerance of the organism. For this reason, the redesing of current biological and physical therapies or the conception of new anticancer therapies aimed at the host. In both cases, these therapies have to be safe, induce minimal adverse events in the host, effective and low cost (compared to existing anticancer therapies).

A second strategy is the stimulation of the action of a continuous periodic external perturbation. Under the action of a periodic external perturbation appears a limit cycle, i.e., both populations oscillate and may coexist indefinitely without extinction. This result is predicted very early by ecological model and more recent it is found in the study of dynamical models of the immune system.

It results interesting that for higher frequencies of the periodic external perturbation, the limit cycle becomes a non-chaotic attractor, which reduces its size as the frequency of this perturbation is increased. This behavior is also observed in the three-dimensional model. These results indicate that malignant tumors may be controlled, but not cured. The latter is of great importance and relevance in cancer therapy since this disease may become chronically controlled (stationary partial response [27]). A merit of this study is that it demonstrates that cancer may be controlled through the appropiate application of a periodic external perturbation to the host and not only to the tumor. The same idea of cancer control, instead of its complete cure, has been reported by other authors from the approach of evolutionary ecology [39, 79–81].

Undoubtedly, these theoretical results need to be tested in preclinical studies and subsequently in clinical ones. This will allow us to evaluate the validity, feasibility and accuracy of the model proposed in this study, as well as the search for others new appropiate periodic external perturbations to the host. Consequently, the LV model may serve as a guide for the design and interpretation of experiments.

The results obtained for the different values of the frequency for the periodic external perturbation may suggest the possible existence of one or several resonance frequencies that potentiate and harmonize the host that may lead to the host governing indefinitely the tumor-host dynamics or the tumor complete remission. This aspect can be addressed in an additional study, taking into account the results reported in previous studies [82–84].

Finally, it is important to point out that the behavior of the higher dimensional LV model, taking into account the effect of other types of cells, is a problem that require further studies. Not only because they are mathematically interesting, but also because they may be crucial in the control of the cancer. Furthermore, although the intention of this study is to model the tumor-host dynamics when the latter is perturbed with a constant or periodic external perturbation, it lays the foundations for generalizing the model when anticancer therapies directed ath the tumor and others directed at the host are used, as occur in the combination of immunotherapy with chemotherapy and/or radiotherapy [40–42], and immunotherapy with irreversible electroporation [85–88]. According to our results, periodic external perturbations with different parameters may be used to perturb the tumor and host in order to understand how the tumor-host dynamics changes. Additionally, it is important to take into account the circadian immune system [89]. These aspects will be analyzed in a future study.

## Acknowledgements

The research in this paper was supported by project PID2022-141385NB-I00 granted by Ministerio de economía y competitividad and by project E41 23R granted by the Aragon Government. Dr. Rolando Placeres thanks Professor Keysner Boet (Physics department, Miami Dade College, USA) for his kindness assistance in the correction of the manuscript. Furthermore, Dr. Luis Bergues and Dr. Juan Montijano thank Dr. Rolando Pèrez Rodríguez (Center for Molecular Immunology) for his valuable suggestions.

